# Insight into RNA-DNA primer length counting by human primosome

**DOI:** 10.1101/2022.05.02.490354

**Authors:** Andrey G. Baranovskiy, Alisa E. Lisova, Lucia M. Morstadt, Nigar D. Babayeva, Tahir H. Tahirov

## Abstract

The human primosome, a four-subunit complex of primase and DNA polymerase alpha (Polα), synthesizes chimeric RNA-DNA primers for DNA polymerases delta and epsilon to initiate DNA replication on both chromosome strands. Despite recent structural insights into the action of its two catalytic centers, the mechanism of DNA synthesis termination is still unclear. Here we report results of functional and structural studies revealing how the human primosome counts RNA-DNA primer length and timely terminates DNA elongation. Using a single-turnover primer extension assay, we defined two factors that determine a mature primer length (~35-mer): 1) a tight interaction of the C-terminal domain of the DNA primase large subunit (p58_C_) with the primer 5’-end, and 2) flexible tethering of p58_C_ and the DNA polymerase alpha catalytic core domain (p180_core_) to the primosome platform domain by extended linkers. The obtained data allows us to conclude that p58_C_ is a key regulator of all steps of RNA-DNA primer synthesis. The above-described findings provide a notable insight into the mechanism of DNA synthesis termination by a eukaryotic primosome, an important process for ensuring successful primer handover to replication DNA polymerases and for maintaining genome integrity.

## INTRODUCTION

Human primosome is a multifunctional complex with a key role in DNA replication (1,2). It is also implicated in a variety of other cellular processes, including telomere maintenance (3,4), innate immunity (5–7), and genome stability (8–10), and it is an emerging candidate for anticancer therapy (11). Primosome is the essential enzyme producing RNA-DNA chimeric primers for the replication of both leading and lagging strands after recruiting either DNA polymerase epsilon (Polε) or delta (Polδ) (12,13). The human primosome comprises the two-subunit primase, with catalytic (p49) and regulatory (p58) subunits, and the two-subunit DNA polymerase α (Polα), with catalytic (p180) and accessory (p70) subunits (1). p180 contains a catalytic core (p180_core_) and a C-terminal domain (p180_C_), while p58 has N-terminal (p58_N_) and C-terminal (p58_C_) domains. Primosome forms an elongated platform p49-p58_N_-p180_C_-p70, which holds p58_C_ and p180_core_ either stationary or flexibly by linkers p58_N_–p58_C_ (L1, residues 253-270) and p180_core_–p180_C_ (L2, residues 1250-1267) (6,14–16) (Figure 1).

**Figure 1.**
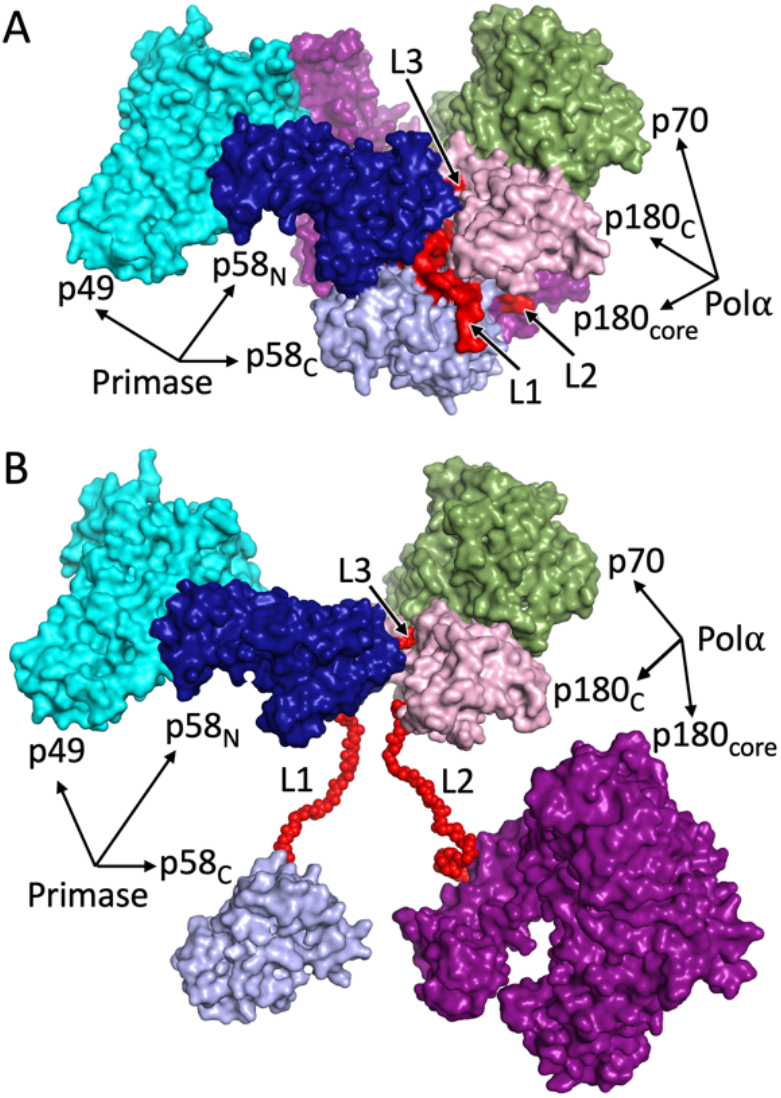
Architecture of human primosome. The platform (p49-p58_N_-p180_C_-p70) can hold p58_C_ and p180_core_ either stationary, as in apo-form (A) or flexibly by linkers (B). The figures were produced using crystal structure of primosome apo-form (PDB accession code 5exr). In panel B the positions of p58_C_ and p180_core_ were moved to arbitrary positions. The linkers L1 and L2 were modeled to show their relative locations. Additional flexibility to platform is provided by linker L3.

The primase is responsible for the initiation, elongation, and termination of RNA primer synthesis when its length reaches nine nucleotides (nt) (15,17–19), and then for its intramolecular transfer to the catalytic subunit of Pola (1,20). After receiving the RNA primer, Pola extends it using deoxyribonucleotide-triphosphates (dNTPs). The concerted RNA-DNA primer synthesis by the two catalytic centers of primosome is tightly regulated; Pola is inactive while primase works, and *vice versa* (15,20). Previously we explored the structural basis for such regulation, including the initiation, elongation, and termination steps of RNA primer synthesis by primase and the following primer transfer to Pola (15). Despite this breakthrough, the factors that regulate the subsequent steps resulting in generation of an RNA-DNA primer of defined length are unknown.

In this work, we conducted a series of biochemical experiments along with structure-based modeling studies to gain insight into the intrinsic mechanism of DNA synthesis elongation and termination by eukaryotic primosome.

## MATERIALS AND METHODS

### Protein expression and purification

Expression and purification to homogeneity of the human primase heterodimer and its mutants (14,21), human primosome (p49-p58-p180-p70) (22), and the Polα catalytic core (22) have been described elsewhere. As previously reported, the N-terminus of p180 (residues 1-334) has been deleted because it is poorly folded (4) and is not required for activity and interaction with other subunits (15). In the deletion mutants Δ5, Δ10, Δ15, and Δp58_C_, the following regions were removed in p58: 256-260, 256-265, 256-270, and 266-560, respectively. In the mutant Ins5, residues GSASG were added after Ser263 of p58. PolαΔ8 with a deleted region 1254-1261 was generated in this work by the site mutagenesis protocol according to (23).

### Oligonucleotides for functional studies

Sequences of all oligonucleotides are provided in Table 1. Oligonucleotides without 5’-triphosphate were obtained from IDT, Inc. The 5-mer RNA primer P5 containing the 5’-triphosphate (5’-pppGGCGG) was obtained as described previously using DNA duplex T8:P4 and the RNA polymerase of bacteriophage T7 (24). Template:primers containing a chimeric RNA-DNA primer P2 were obtained by ligation using the corresponding template, the primers P5 and P6 (Figure 2), and RNA ligase 2 of bacteriophage T4 (New England BioLabs, Inc.). Reactions were incubated for one hour at 25 °C and ligated duplexes were purified by 1ml monoQ column (Cytiva) at 50 °C.

**Figure 2.**
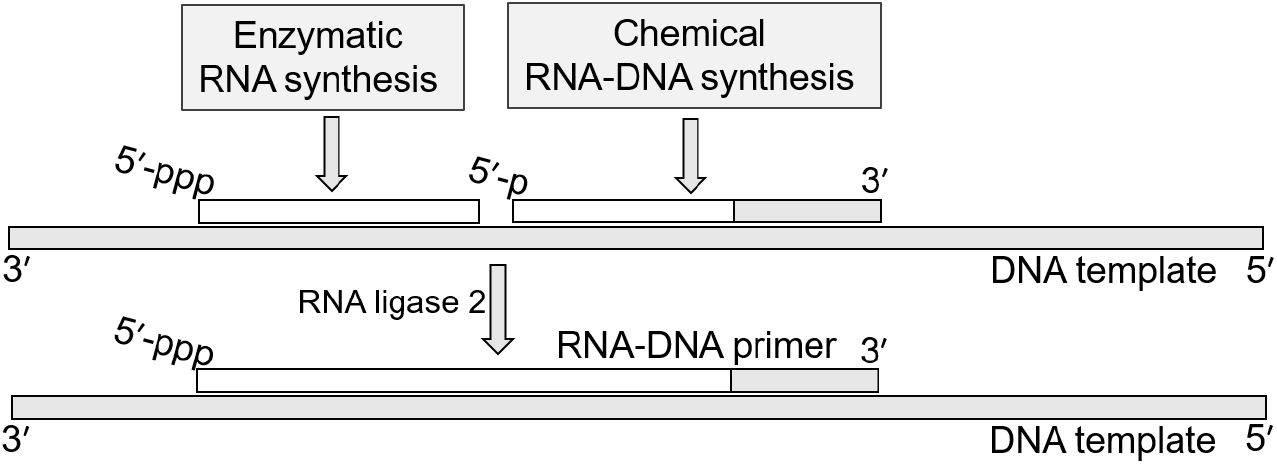
A scheme of 12-mer RNA-DNA primer synthesis. The 5-mer RNA primer with a 5’-triphosphate was generated by RNA polymerase of bacteriophage T7. The 7-mer RNA-DNA with a 5’-phosphate and three dNMPs at the 3’-end was synthesized by IDT Inc. Both primers were annealed to a DNA template and ligated by RNA ligase 2 of bacteriophage T4 (New England BioLabs Inc.). White and gray blocks represent RNA and DNA, respectively.

**Table 1.**
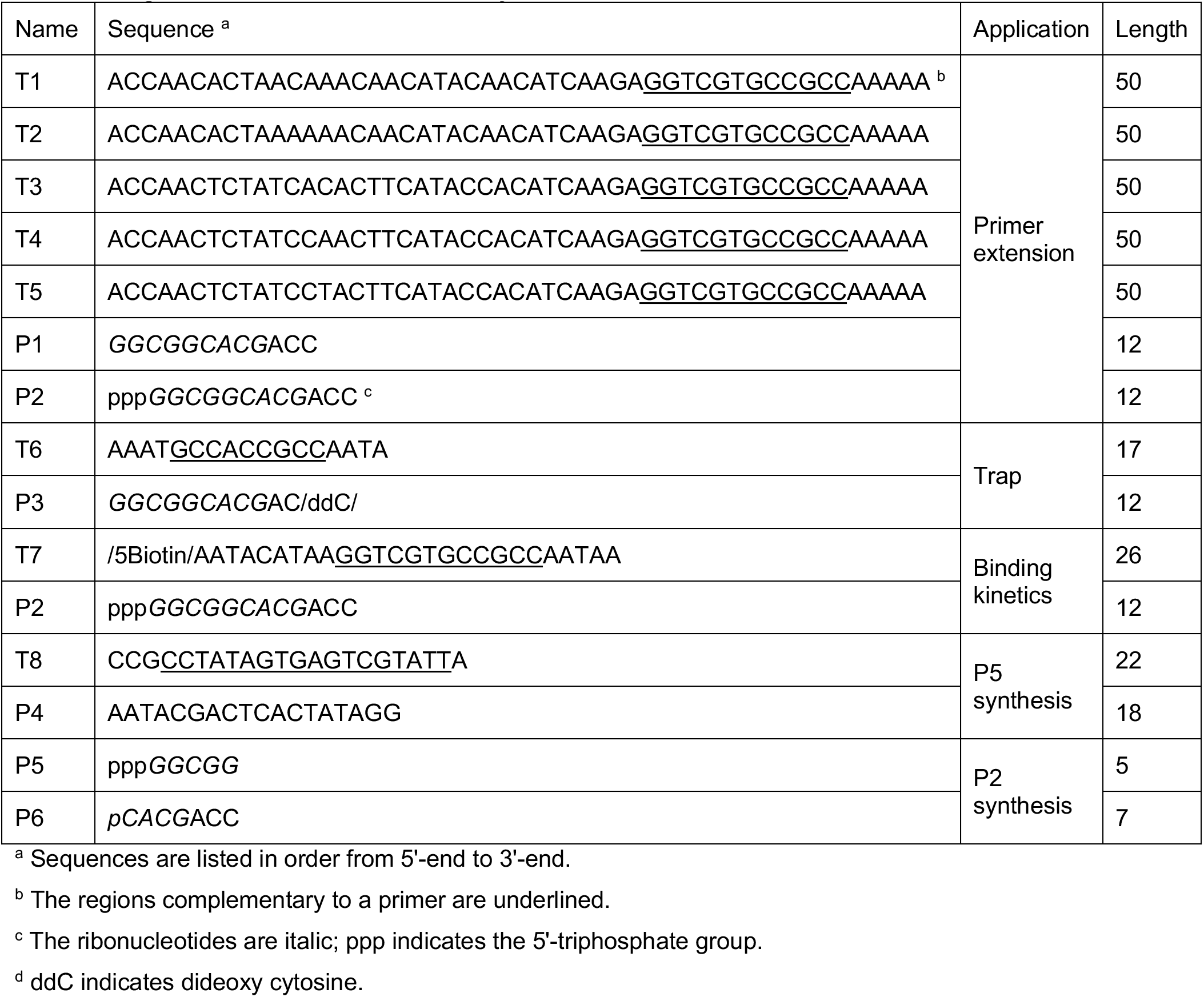
Oligonucleotides used in this study.

### Primer extension assay

DNA-synthetic activity was tested in 10 μl reaction containing 0.6 μM template:primer, 0.2 μM enzyme, 10 μM dNTPs, 0.1 μM [α-^32^P]-dCTP (3000 Ci/mmol; PerkinElmer, Inc.), 10 μM trap, and the buffer: 30 mM Tris-Hepes, pH 7.8, 120 mM KCl, 30 mM NaCl, 1% glycerol, 2 mM TCEP, 5 mM MgCl_2_, and 0.2 mg/ml BSA. The trap was a T6:P3 duplex containing the dideoxy-cytosine at the primer 3’-end to make a dead-end complex with Polα. All primosome mutants were prepared in the reaction buffer by mixing 1 μM Polα or PolαΔ8 with a corresponding primase mutant added at 20% molar excess (Supplemental Figure S1). The enzyme was pre-incubated with a template:primer in 5 μl for 1 min on ice and for 10 sec at 35 °C, then reaction was initiated by addition of 5 μl solution containing dNTPs and trap. Reactions were incubated in PCR tubes on a water bath for 30 sec at 35 °C and stopped by mixing with equal volume of formamide loading buffer (90% v/v formamide, 50 mM EDTA, pH 8, 0.02% Bromophenol blue), heated at 95 °C for 1 min, and resolved by 20% Urea-PAGE (UreaGel System (19:1 Acrylamide/Bisacrylamide), National Diagnostics) for 2.5 h at 3000 V. The reaction products were visualized by phosphorimaging (Typhoon FLA 9500, Cytiva). All activity gels were repeated at least two times.

### Binding studies

Analysis of binding kinetics was done at 23 °C on Octet K2 (Sartorius AG). A template T7 with a biotin at the 5’-end annealed to the primer P2 (Table 1) was prepared by ligation and immobilized on a streptavidin-coated biosensor (SAX, Sartorius AG) at 50 nM concentration for 7 min at 500 rpm; sensors were then blocked by incubating for 2 min in 10 μg/ml biocytin. Binding studies were conducted in a 96-well microplate (Greiner Bio-One) in buffer containing 30 mM Tris-Hepes, pH 7.8, 150 mM NaCl, 5 mM MgCl2, 2 mM TCEP, and 0.002% Tween 20. Each binding cycle starts with a baseline step by incubating a sensor for 30 sec in a well with buffer. After that, the sensor moves to a well with p58_C_ solution to begin an association step. After sensor saturation with an analyte (p58_C_), it returns to the baseline well to start a dissociation step. Approximately 20 min was required for complete dissociation of p58_C_ from the sensor loaded with T7:P2. All steps were conducted at shaking speed of 1,000 rpm. Data Analysis HT software (ver. 11.1, Sartorius AG) was used to calculate binding constants (*k_on_, k_off_*, and *K_D_*) by using global fitting. The average value and standard deviation were calculated from three independent experiments.

### Modeling

For the modeling of primosome elongation complexes, we used the coordinates of primosome in apo form (pdb code 5exr) (15), p180_core_ with an RNA-primed DNA template and dCTP (4qcl) (25), p58_C_ with an RNA-primed DNA template (5f0q) (15), and an ideal B-DNA generated with Coot (26). Modeling was performed by constructing several template:primer duplexes up to 33-base-pair (bp) by combining the duplex parts from the crystal structures as well as ideal B-DNA in the case of duplexes over 15-bp, by manual adjustment, and by using “regularize zone” option of Coot software (26).

## RESULTS

### Primosome autonomously controls the length of an RNA-DNA primer

The design of our experiments was based on the hypothesis that p58_C_ stays bound to the template:primer upon RNA primer transfer from p49 to Polα and the following primer extension with dNTPs. This hypothesis is based on the high affinity of p58_C_ to the template:primer (*K*_d_ = 36 nM) (24) and its ability to share the 9-mer RNA primer with p180_core_ (1,15). We first compared primer extension by p180_core_ and primosome using a pre-annealed template:primer, which mimics the native primosome substrate (1). It presents a 50-mer DNA template annealed to a 12-mer chimeric primer (5’-pppGGCGGCACG<u>ACC</u>, DNA region is underlined) containing nine ribonucleotides with the triphosphate group at the 5’-end and three deoxynucleoside monophosphates (dNMPs) at the 3’-end (Figure 2). To observe the products generated during a single round of primer extension, a DNA trap (T6:P3 template:primer with the blocked 3’-end; Table 1) was added together with dNTPs at reaction start to prevent repriming events. For example, in the absence of a DNA trap, p180_core_ mainly generates a 45-mer product, which suggests multiple binding events (Figure 3A, lane 1). At single-turnover conditions (with a DNA trap), we observe the actual p180_core_ processivity (lane 2), which is significantly lower than in the absence of a trap.

**Figure 3.**
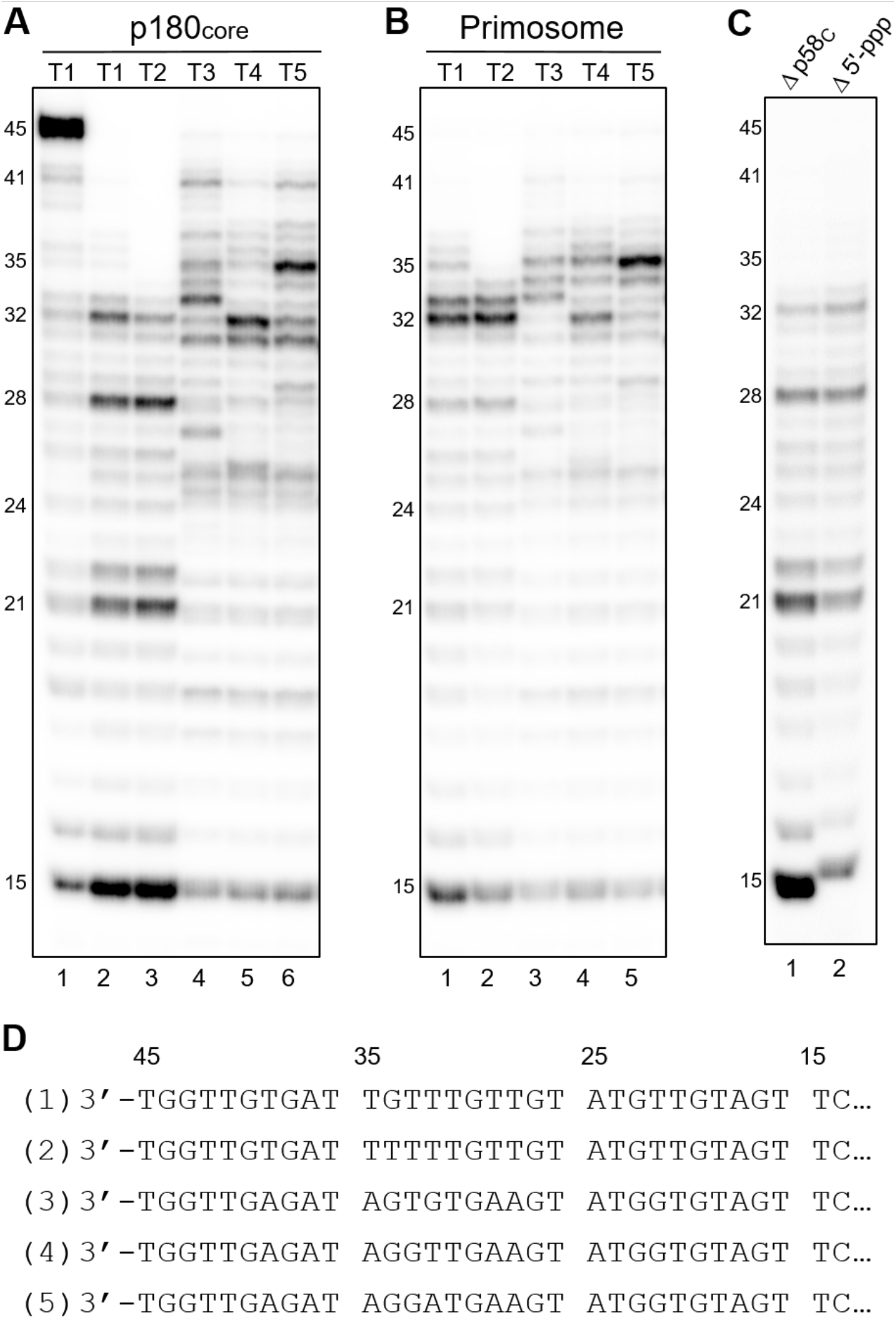
Analysis of DNA synthesis elongation and termination by p180_core_ and primosome. The products generated by p180_core_ and primosome on templates T1-T5 annealed to the 12-mer primer P2 are shown on panels **(A)** and **(B)**, respectively. **(C)** Primosome shows reduced processivity of DNA synthesis when the primer-p58_C_ interaction is disrupted. Lane 1 – T1:P2 and primosome with deleted p58_C_; lane 2 – T1:P1 (primer without 5’-triphosphate) and primosome. **(D)** Sequences of synthesized primers. The numbers of corresponding templates are shown in commas. Reactions contained 0.6 μM template:primer, 0.2 μM enzyme, 10 μM dNTPs, 0.1 μM [α-^32^P]-dCTP, and 10 μM trap (except lane 1 on panel A), and were incubated at 35 °C for 30 sec. The products were resolved by 20% Urea-PAGE and visualized by phosphorimaging.

A comparison of the primer extension products generated at single-turnover conditions on the templates T1 and T2 by p180_core_ (Figure 3A, lanes 2 and 3) and primosome (Figure 3B, lanes 1 and 2) revealed that primosome makes longer products, indicative of increased processivity of DNA synthesis. On the templates with a reduced AT-content, p180_core_ showed higher processivity of DNA synthesis than on T1 and T2 (Figure 3A, lanes 4-6, compared to lanes 2 and 3) resulting in generation of a notable amount of 35-41-mer products. In contrast, when the primosome performed DNA synthesis on T3-T5, the level of primers with a length exceeding 37nt is significantly reduced (Figure 3B, lanes 3-5, compared to Figure 3A, lanes 4-6). Thus, Polα processivity significantly depends on the template sequence, while for primosome such dependence is much weaker.

These data indicate that primosome has the ability to regulate primer elongation by Polα and to terminate DNA synthesis, which results in generation of RNA-DNA primers with a length of 32-37nt, defined here as mature primers. Products with a similar length were obtained in the course of pulse-chase experiments conducted in the SV40 replication system (27). Our data revealed that besides Polα there are additional factors in primosome that control the primer length. These factors increase processivity of DNA synthesis when the primer is shorter than 32-mer and gradually reduce processivity when its length exceeds 37nt.

### p58_C_ and primosome linkers define the length of an RNA-DNA primer

To test the hypothesis that p58_C_ regulates the primer length by holding the primer 5’-end (24), we repeated the assay using a primosome with deleted p58_C_ or a template:primer without a 5’-triphosphate on the primer strand. In both cases the primosome generated shorter products (Figure 3C, lanes 1 and 2, compared to lane 1 of panel B), similar to that seen in reactions with p180_core_ (Figure 3A, lane 2). Overall, these results indicate that p58_C_ is associated with the primer 5’-end when p180_core_ extends the primer 3’-end. Thus, in primosome, p58_C_ works as a processivity factor for Polα, providing quick intramolecular reloading of the template:primer to the active center of a DNA polymerase. This conclusion is supported by the fact that the DNA trap is not efficient in blocking the Polα active site during synthesis of a 32-mer primer by primosome (Figure 3B).

The two-point interaction mode between the primer and primosome, with p58_C_ at the 5’-end and p180_core_ at the 3’-end, predicts an increase in the distance between these domains during primer extension and stretching of the linkers L1 and L2, which tether p58_C_ and p180_core_ to the platform. To test this hypothesis, we generated one Polα mutant by deleting 8aa in L2 as well as four primase mutants by insertion of five amino acids (aa) or by deleting five, ten, and fifteen aa in L1. Eight primosome mutants, containing different combinations of linkers in primase and Polα, were obtained (Supplemental Figure S1) and tested in a primer extension assay on T1 (Figure 4). The 5aa extension of L1 resulted in 1.9-fold increase in the level of 35- to 37-mer products versus those 32- to 33-mer in length (Figure 4A, comparison of lanes 1 and 3, and Figure 4B). L1 shortening by 5aa had a small effect on products distribution (Figure 4A, lane 5), while deletion of 10aa and especially 15aa significantly raised the level of immature primers (lanes 7-10). For example, the level of a 28-mer product increased 3.4-fold in the case of a 15aa deletion in L1 and 7-fold when this L1 truncation was combined with an 8aa deletion in L2 (Figure 4B).

**Figure 4.**
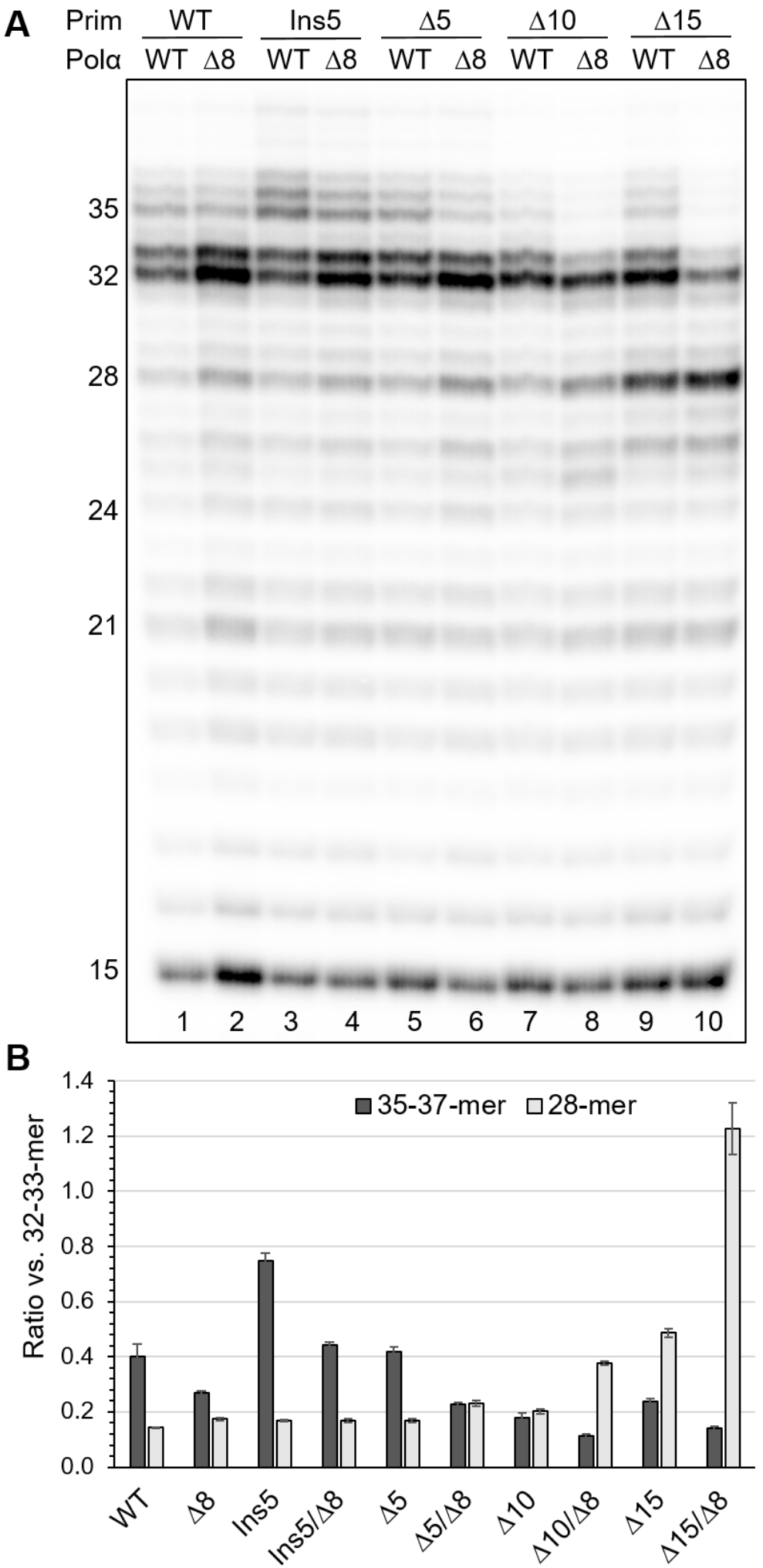
The linkers in primase and Polα regulate the primer length. (**A**) Analysis of primer extension products generated by primosome and its mutants. Lane 1 – T1:P2 and primosome with intact linkers. Lanes 2-10 – T1:P2 and primosome mutants with linker modifications in primase and Polα. Δ5, Δ10, Δ15, and Ins5 denotes deletion of 5aa, 10aa, 15aa, and insertion of 5aa into primase linker, respectively. Δ8 denotes deletion of 8aa in Polα linker. (**B**) Effect of linkers length on the level of selected products (see the panel A). The data are presented as bar graphs showing the mean ±SD calculated from three independent experiments (Supplemental Figure S2).

In summary, these data support the idea that the L1 and L2 linkers play a role in DNA synthesis termination by controlling the primer length. Of note, Polα residues 1240-1249 may turn to the coil (Figure 5) and become the part of L2 extending its length to 28aa. Moreover, there is a L3 linker in p180 (1445–1447) that tethers p180_C_ to p58_N_ and makes the platform flexible between the points of L1 and L2 attachment to it (Figure 1). These factors may help the mutated primosome to partially compensate for a reduction in linker length.

**Figure 5.**
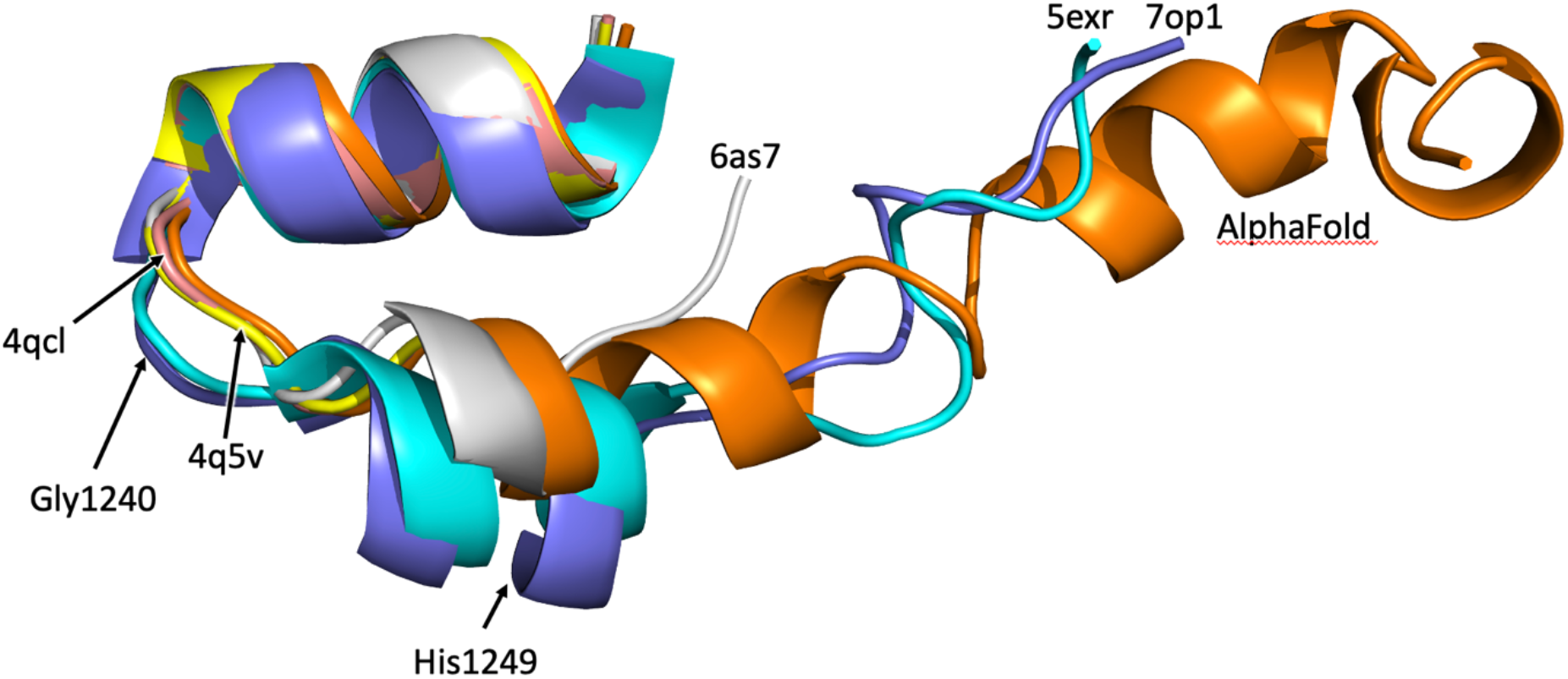
Comparison of Pola structures revealing different conformations or partial disorder of p180 residues 1240-1249. PDB accession numbers of used coordinates are indicated. The L2 structure predicted by AlphaFold is also used for alignment.

### Stability of p58_C_/template:primer complex defines DNA synthesis termination

We estimated stability of the p58_C_ complex with a T7:P2 duplex containing the 12-mer chimeric RNA-DNA primer that was used in primer extension studies (Table 1) and has a 5’-triphosphate. Binding studies were conducted on Octet K2, which allows for extracting the rate constants of complex formation (*k*_on_) and dissociation (*k*_off_) and calculating the dissociation constant (*K_D_*). Recently, we applied this approach to studying the binding kinetics for Polα and DNA (28). A 26-mer DNA template T7 with biotin at the 5’-end was primed by a 12-mer primer P2 and loaded on a streptavidin-coated sensor.

An obtained *k*_off_ value of 0.004 sec^-1^ points out to a stable p58_C_/T7:P2 complex with a half-life of ~3 min (Table 2). In comparison, Polα shows 11-fold and 85-fold weaker interaction with a template:primer in the presence and absence of incoming dNTP, respectively. According to these data, Polα has a significantly higher probability than p58_C_ to dissociate from the mature primer when the tension in the stretched linkers is built up, resulting in DNA synthesis termination. The *K_D_* value of 14 nM obtained here for the p58_C_/T7:P2 complex (Table 2) is 2-fold lower than the value that we obtained previously by using a gel-retardation assay and a 7-mer primer (24). Taking into account the differences in employed approaches and experimental setup, we can conclude that in general these results are consistent.

**Table 2.**
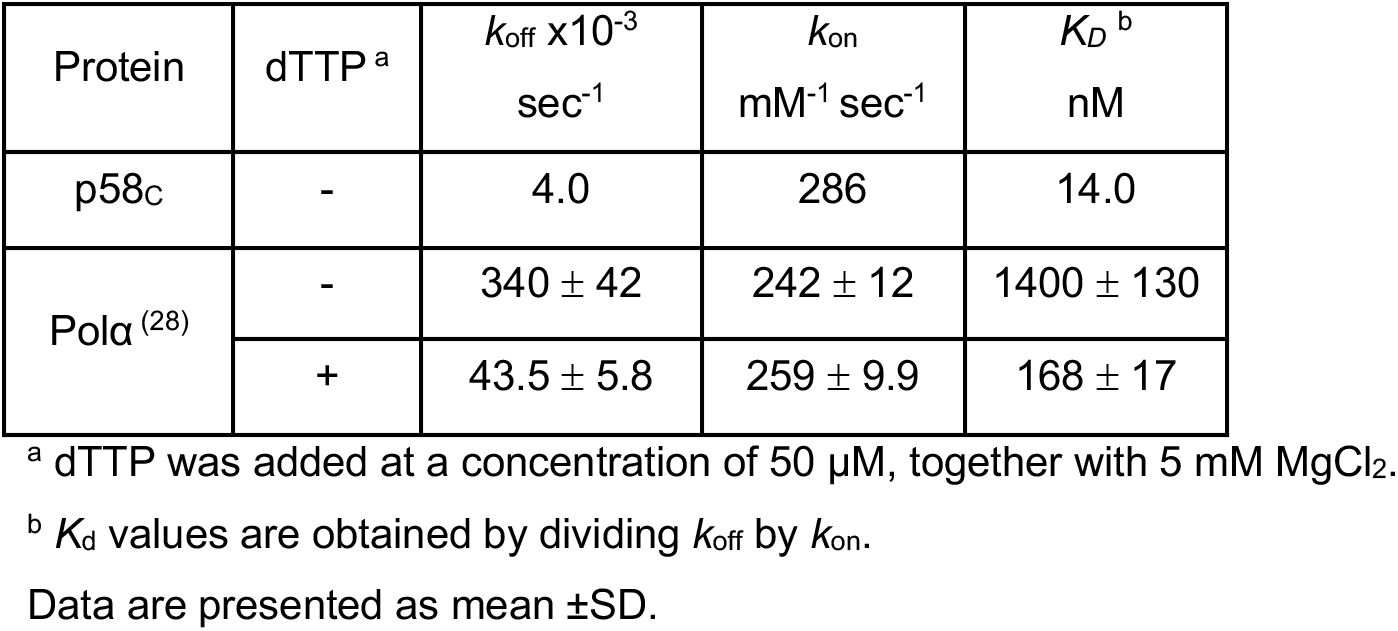
Results of binding studies.

To evaluate the consistency of our data with the structural potential of primosome to stretch the linkers, we obtained several structure-based models. At initiation of dNMPs incorporation by Pola, the linker length is sufficient for free spiral rotational movement of p58_C_ relative to p180_core_. However, the options for linkers become increasingly limited during the last helical turn. As illustrated in Figure 6A with a 23-mer RNA-DNA primer, both the L1 and L2 linkers are in extended conformation, but still not fully stretched. Addition of a half helical turn to the template-primer duplex moves distant points of linkers closer to the platform, thus providing some relaxation for the linkers. Upon addition of more dNMPs, these ends of L1 and L2 move away from the platform, resulting in almost fully stretched linkers with 32- or 33-mer primers (Figure 6B).

**Figure 6.**
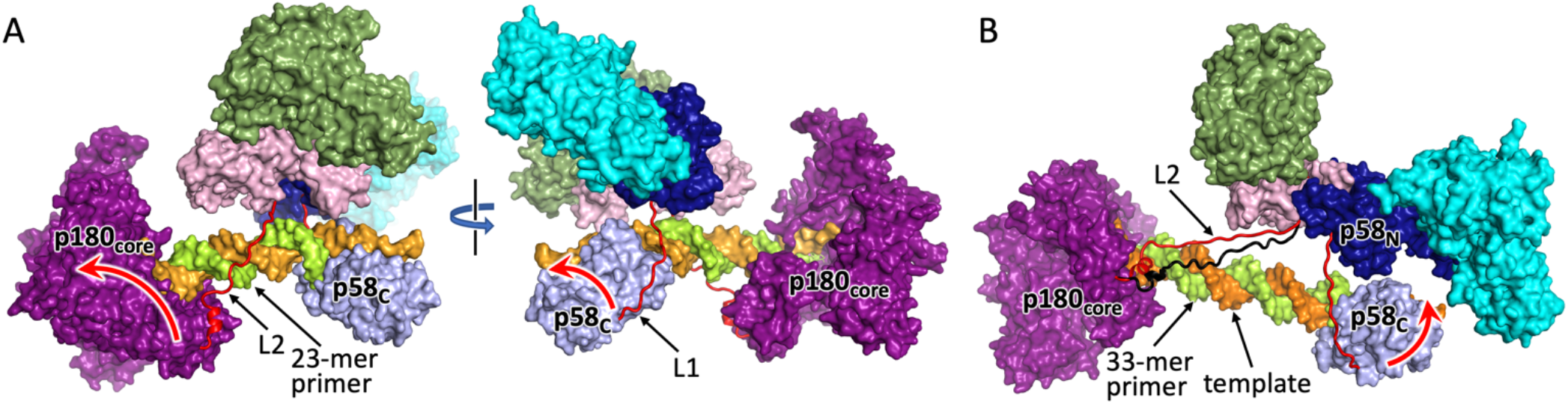
Structure-based models of primosome elongation complexes. Models with 23-mer (**A**) and 33-mer primers (**B**) are shown. Coloring of domains is the same as in Figure 1. The linkers are represented as cartoons and colored red. Red arrows indicate the direction of domain movement upon primer extension. In panel (**B**), L2 with a partially unfolded helix is colored black.

## DISCUSSION

Our studies unveiled the intricate mechanism of RNA-DNA primer length counting by primosome. This mechanism is based on stable p58_C_ interaction with a template:primer, which boasts Polα processivity during primer maturation. The two-point binding mode of a template:primer by primosome, with p58_C_ at the primer 5’-end and p180_core_ at the 3’-end, dictates the involvement of primosome linkers L1 and L2 in primer length counting and DNA synthesis termination (Figure 6). These linkers, with a cumulative length of 36aa, act as springs, requiring some energy for stretching. Probably, this energy comes from hydrolysis of the phosphodiester bond of dNTPs. Upon addition of 23-28 dNMPs, the accumulated tension in the stretched linkers gradually increases probability of p180_core_ ejection from the duplex and DNA synthesis termination. Once the mature primer is dissociated from p180_core_, reloading of p58_C_-bound template:primer to the Polα active site is complicated because it requires significant spontaneous stretching of the linkers. Thus, p58_C_ is a crucial element in DNA synthesis termination. Due to the mutagenic potential of Polα (8), which does not possess proofreading exonuclease activity, limitation of a DNA track to ~25nt is important for genome stability.

The obtained data allow us to conclude that p58_C_ is a critical regulator of all steps of RNA-DNA primer synthesis, from RNA synthesis initiation by primase until DNA synthesis termination by Polα. After initiation, elongation, and termination of RNA-DNA primer synthesis, p58_C_ may stay bound to the 5’-end of a primer and, consequently, participates in primer handoff from Polα to Polε and Polδ, possibly by facilitating RFC/PCNA loading. Interestingly, because p58_C_ is also responsible for initiation of RNA primer synthesis, primosome stays autoinhibited until p58_C_ dissociates from the previously synthesized primer. Considering the relatively long half-life of the complex with a template:primer, p58_C_ may remain bound to it until completion of Okazaki fragment synthesis that only takes seconds (29,30). Thus, p58_C_ recycling may emerge as a key aspect of controlling events at the replication fork, both spatially and temporally. For example, p58_C_ may prevent PCNA from sliding off the primer 5’-end after RFC dissociation.

## DATA AVIABILITY

The data that support the findings of this study are included in the Supplementary Data file or available from the corresponding author upon request.

## ACKNOWLEDGEMENT

We thank K. Jordan for editing this manuscript.

## FUNDING

This work was supported by the National Institute of General Medical Sciences grant R35GM127085 to T.H.T.

### Conflict of interest statement

None declared

## Author Contributions

A.G.B developed the protocol for chimeric primer synthesis, carried out the biochemical experiments, and supervised preparation of samples for functional studies. L.M.M, A.E.L, and N.D.B participated in preparation of samples and technical support. T.H.T initiated the project, performed modeling, and proposed and clarified the mechanism of termination with the help of team members. T.H.T and A.G.B wrote the manuscript.

## Notes

### Competing Interest Statement

The authors have declared no competing interest.

## REFERENCES

1. Baranovskiy, A.G. and Tahirov, T.H. (2017) Elaborated Action of the Human Primosome. Genes (Basel), 8.

2. Pellegrini, L. (2012) The Pol alpha-primase complex. Subcell Biochem, 62, 157–169.

3. Casteel, D.E., Zhuang, S., Zeng, Y., Perrino, F.W., Boss, G.R., Goulian, M. and Pilz, R.B. (2009) A DNA polymerase-{alpha}{middle dot}primase cofactor with homology to replication protein A-32 regulates DNA replication in mammalian cells. J Biol Chem, 284, 5807–5818.

4. Mizuno, T., Hirabayashi, K., Miyazawa, S., Kobayashi, Y., Shoji, K., Kobayashi, M., Hanaoka, F., Imamoto, N. and Torigoe, H. (2021) The intrinsically disordered N-terminal region of mouse DNA polymerase alpha mediates its interaction with POT1a/b at telomeres. Genes Cells, 26, 360–380.

5. Starokadomskyy, P., Gemelli, T., Rios, J.J., Xing, C., Wang, R.C., Li, H., Pokatayev, V., Dozmorov, I., Khan, S., Miyata, N. et al. (2016) DNA polymerase-alpha regulates the activation of type I interferons through cytosolic RNA:DNA synthesis. Nat Immunol, 17, 495–504.

6. Kilkenny, M.L., Veale, C.E., Guppy, A., Hardwick, S.W., Chirgadze, D.Y., Rzechorzek, N.J., Maman, J.D. and Pellegrini, L. (2022) Structural basis for the interaction of SARS-CoV-2 virulence factor nsp1 with DNA polymerase alpha-primase. Protein Sci, 31, 333–344.

7. Tang, L., Sheraz, M., McGrane, M., Chang, J. and Guo, J.T. (2019) DNA Polymerase alpha is essential for intracellular amplification of hepatitis B virus covalently closed circular DNA. PLoS Pathog, 15, e1007742.

8. Reijns, M.A.M., Kemp, H., Ding, J., de Proce, S.M., Jackson, A.P. and Taylor, M.S. (2015) Lagging-strand replication shapes the mutational landscape of the genome. Nature, 518, 502–506.

9. Schimmel, J., Munoz-Subirana, N., Kool, H., van Schendel, R. and Tijsterman, M. (2021) Small tandem DNA duplications result from CST-guided Pol alpha-primase action at DNA break termini. Nat Commun, 12, 4843.

10. Mirman, Z., Lottersberger, F., Takai, H., Kibe, T., Gong, Y., Takai, K., Bianchi, A., Zimmermann, M., Durocher, D. and de Lange, T. (2018) 53BP1-RIF1-shieldin counteracts DSB resection through CST- and Polalpha-dependent fill-in. Nature, 560, 112–116.

11. Han, T., Goralski, M., Capota, E., Padrick, S.B., Kim, J., Xie, Y. and Nijhawan, D. (2016) The antitumor toxin CD437 is a direct inhibitor of DNA polymerase alpha. Nat Chem Biol, 12, 511–515.

12. Burgers, P.M.J. and Kunkel, T.A. (2017) Eukaryotic DNA Replication Fork. Annu Rev Biochem, 86, 417–438.

13. Pellegrini, L. and Costa, A. (2016) New Insights into the Mechanism of DNA Duplication by the Eukaryotic Replisome. Trends Biochem Sci, 41, 859–871.

14. Baranovskiy, A.G., Zhang, Y., Suwa, Y., Babayeva, N.D., Gu, J., Pavlov, Y.I. and Tahirov, T.H. (2015) Crystal structure of the human primase. J Biol Chem, 290, 5635–5646.

15. Baranovskiy, A.G., Babayeva, N.D., Zhang, Y., Gu, J., Suwa, Y., Pavlov, Y.I. and Tahirov, T.H. (2016) Mechanism of Concerted RNA-DNA Primer Synthesis by the Human Primosome. J Biol Chem, 291, 10006–10020.

16. Nunez-Ramirez, R., Klinge, S., Sauguet, L., Melero, R., Recuero-Checa, M.A., Kilkenny, M., Perera, R.L., Garcia-Alvarez, B., Hall, R.J., Nogales, E. et al. (2011) Flexible tethering of primase and DNA Pol alpha in the eukaryotic primosome. Nucleic Acids Res, 39, 8187–8199.

17. Sheaff, R.J. and Kuchta, R.D. (1993) Mechanism of calf thymus DNA primase: slow initiation, rapid polymerization, and intelligent termination. Biochemistry, 32, 3027–3037.

18. Copeland, W.C. and Wang, T.S. (1993) Enzymatic characterization of the individual mammalian primase subunits reveals a biphasic mechanism for initiation of DNA replication. J Biol Chem, 268, 26179–26189.

19. Kuchta, R.D., Reid, B. and Chang, L.M. (1990) DNA primase. Processivity and the primase to polymerase alpha activity switch. J Biol Chem, 265, 16158–16165.

20. Sheaff, R.J., Kuchta, R.D. and Ilsley, D. (1994) Calf thymus DNA polymerase alpha-primase: “communication” and primer-template movement between the two active sites. Biochemistry, 33, 2247–2254.

21. Baranovskiy, A.G., Gu, J., Babayeva, N.D., Agarkar, V.B., Suwa, Y. and Tahirov, T.H. (2014) Crystallization and preliminary X-ray diffraction analysis of human DNA primase. Acta Crystallogr F Struct Biol Commun, 70, 206–210.

22. Zhang, Y., Baranovskiy, A.G., Tahirov, T.H. and Pavlov, Y.I. (2014) The C-terminal domain of the DNA polymerase catalytic subunit regulates the primase and polymerase activities of the human DNA polymerase alpha-primase complex. J Biol Chem, 289, 22021–22034.

23. Liu, H. and Naismith, J.H. (2008) An efficient one-step site-directed deletion, insertion, single and multiple-site plasmid mutagenesis protocol. BMC Biotechnol, 8, 91.

24. Baranovskiy, A.G., Zhang, Y., Suwa, Y., Gu, J., Babayeva, N.D., Pavlov, Y.I. and Tahirov, T.H. (2016) Insight into the Human DNA Primase Interaction with Template-Primer. J Biol Chem, 291, 4793–4802.

25. Baranovskiy, A.G., Duong, V.N., Babayeva, N.D., Zhang, Y., Pavlov, Y.I., Anderson, K.S. and Tahirov, T.H. (2018) Activity and fidelity of human DNA polymerase alpha depend on primer structure. J Biol Chem, 293, 6824–6843.

26. Emsley, P., Lohkamp, B., Scott, W.G. and Cowtan, K. (2010) Features and development of Coot. Acta Crystallogr D Biol Crystallogr, 66, 486–501.

27. Murakami, Y., Eki, T. and Hurwitz, J. (1992) Studies on the initiation of simian virus 40 replication in vitro: RNA primer synthesis and its elongation. Proc Natl Acad Sci U S A, 89, 952–956.

28. Baranovskiy, A.G., Babayeva, N.D., Lisova, A.E., Morstadt, L.M. and Tahirov, T.H. (2022) Structural and functional insight into mismatch extension by human DNA polymerase α. Proc Natl Acad Sci U S A, in press.

29. Raghuraman, M.K., Winzeler, E.A., Collingwood, D., Hunt, S., Wodicka, L., Conway, A., Lockhart, D.J., Davis, R.W., Brewer, B.J. and Fangman, W.L. (2001) Replication dynamics of the yeast genome. Science, 294, 115–121.

30. Stodola, J.L. and Burgers, P.M. (2016) Resolving individual steps of Okazaki-fragment maturation at a millisecond timescale. Nat Struct Mol Biol, 23, 402–408.

