## Supplemental Figures 1 and 2 for "Insight into RNA-DNA primer length counting by human primosome"

**for the article**

[1]Eppley Institute for Research in Cancer and Allied Diseases, Fred & Pamela Buffett Cancer Center. University of Nebraska Medical Center, Omaha, NE, USA.

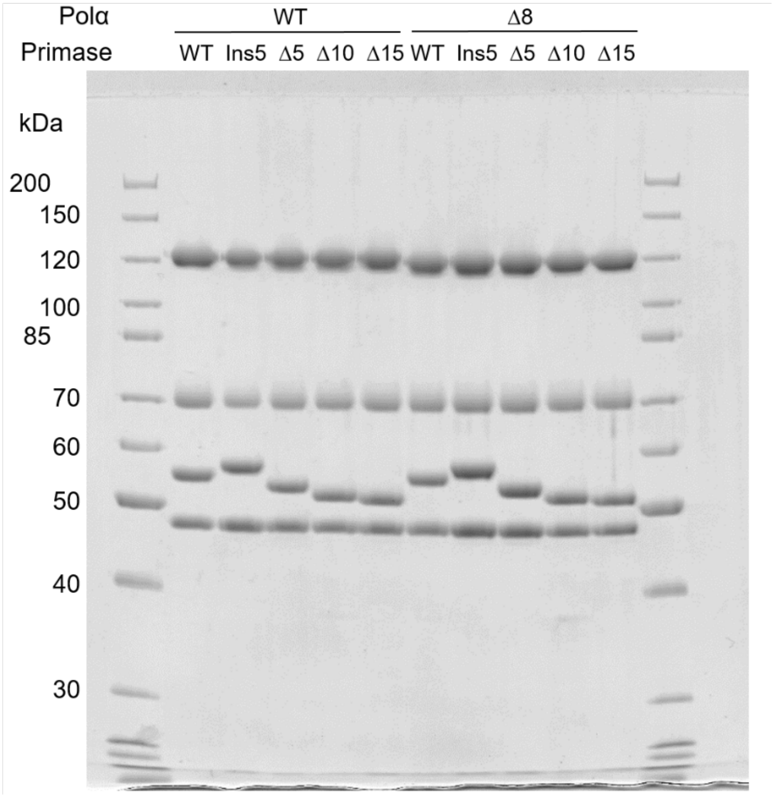


**Supplemental Figure 1. The purity of primosome and its mutants.** Proteins were separated by 8% SDS-PAGE and stained by Coomassie Brilliant Blue R-250. Modifications in linkers of p58 (L1) and p180 (L2) are shown above the wells. A disordered N-terminus of p180 (residues 1-334) is deleted.

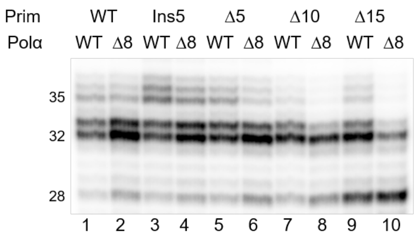


| Product length | Integrated density | | | | | | | | | |
|---|---|---|---|---|---|---|---|---|---|---|
| | lane 1 | lane 2 | lane 3 | lane 4 | lane 5 | lane 6 | lane 7 | lane 8 | lane 9 | lane 10 |
| 35-37 (A) | 6217 | 5262 | 10678 | 9815 | 7864 | 4701 | 2695 | 1546 | 4388 | 1276 |
| 32-33 (B) | 13862 | 19207 | 13810 | 21743 | 18423 | 20893 | 13843 | 13032 | 17673 | 8644 |
| 28 (C ) | 2010 | 3211 | 2267 | 3461 | 2995 | 4609 | 2693 | 4845 | 8451 | 10759 |
| ratio A/B | 0.45 | 0.27 | 0.77 | 0.45 | 0.43 | 0.23 | 0.19 | 0.12 | 0.25 | 0.15 |
| ratio C/B | 0.15 | 0.17 | 0.16 | 0.16 | 0.16 | 0.22 | 0.19 | 0.37 | 0.48 | 1.24 |

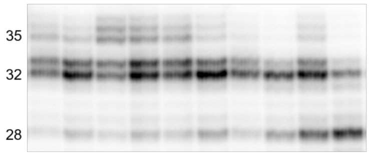


| Product length | Integrated density | | | | | | | | | |
|---|---|---|---|---|---|---|---|---|---|---|
| | lane 1 | lane 2 | lane 3 | lane 4 | lane 5 | lane 6 | lane 7 | lane 8 | lane 9 | lane 10 |
| 35-37 (A) | 5917 | 6474 | 12770 | 9780 | 7752 | 4755 | 2867 | 1439 | 4433 | 1223 |
| 32-33 (B) | 14989 | 24741 | 17816 | 22921 | 19561 | 21889 | 17729 | 14175 | 19345 | 8922 |
| 28 (C ) | 2112 | 4366 | 3012 | 3942 | 3342 | 5226 | 3557 | 5436 | 9725 | 11680 |
| ratio A/B | 0.39 | 0.26 | 0.72 | 0.43 | 0.40 | 0.22 | 0.16 | 0.10 | 0.23 | 0.14 |
| ratio C/B | 0.14 | 0.18 | 0.17 | 0.17 | 0.17 | 0.24 | 0.20 | 0.38 | 0.50 | 1.31 |

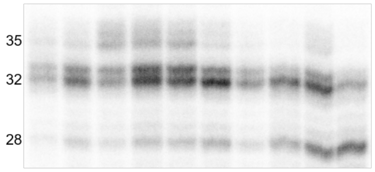


| Product length | Integrated density | | | | | | | | | |
|---|---|---|---|---|---|---|---|---|---|---|
| | lane 1 | lane 2 | lane 3 | lane 4 | lane 5 | lane 6 | lane 7 | lane 8 | lane 9 | lane 10 |
| 35-37 (A) | 2296 | 3098 | 6915 | 7843 | 6976 | 3723 | 1548 | 1262 | 3311 | 932 |
| 32-33 (B) | 6299 | 11375 | 9246 | 17632 | 16335 | 15743 | 8388 | 10801 | 13791 | 6855 |
| 28 (C ) | 914 | 2032 | 1574 | 3046 | 2886 | 3717 | 1763 | 4056 | 6575 | 7697 |
| ratio A/B | 0.36 | 0.27 | 0.75 | 0.44 | 0.43 | 0.24 | 0.18 | 0.12 | 0.24 | 0.14 |
| ratio C/B | 0.15 | 0.18 | 0.17 | 0.17 | 0.18 | 0.24 | 0.21 | 0.38 | 0.48 | 1.12 |

**Supplemental Figure 2. Quantification of gels corresponding to Fig. 4A.** The integrated densities of selected bands were quantified using ImageJ software.
